# Social Context Suppresses Food Anticipatory Activity and Associated Thermoregulation in Mice

**DOI:** 10.64898/2026.04.10.717725

**Authors:** Audrey I. Paik, Jacqueline R. Trzeciak, Cameron Harrington, Andrew D. Steele

## Abstract

Food anticipatory activity (FAA) is a robust behavioral output of food-entrained circadian rhythms, characterized by increased locomotor activity prior to scheduled feeding. Despite the social nature of rodents, FAA is almost exclusively studied in singly housed animals, leaving the influence of social context largely unexplored. Here, we used implanted wireless devices to measure individual locomotor activity and subcutaneous body temperature in group-housed mice and compared these measures to singly housed controls. Social housing significantly suppressed FAA in both male and female mice, with the effect being more pronounced and sustained in males. In parallel, preprandial increases in body temperature were markedly attenuated in group-housed animals. These findings demonstrate that FAA is a flexible, state-dependent behavior that reflects both circadian timing and energetic demand. Together, these results identify social context as an important and underappreciated determinant of food-entrained circadian biology.

## Introduction

Circadian rhythms are intrinsic, self-sustaining oscillations with a period of approximately 24 hours that regulate diverse physiological and behavioral processes, including sleep, temperature regulation, metabolism, and activity [1, 2]. These rhythms are synchronized to the external environment by zeitgebers, or time cues, the most prominent of which is the light–dark (LD) cycle. However, food availability is also a potent zeitgeber, capable of entraining circadian rhythms in both central and peripheral tissues, particularly under time-restricted feeding (TRF) or timed, calorie-restricted (CR) conditions [3, 4].

One of the most striking behavioral consequences of food-based entrainment is food anticipatory activity (FAA), a circadian-controlled state characterized by a strong increase in locomotor activity and a synchronized pre-prandial rise in core body temperature that occurs in the hours preceding scheduled feeding [5, 6]. FAA is the output of the food-entrainable oscillator (FEO), a timing system that is traditionally studied by feeding mice during the daytime when they are normally inactive [7, 8]. Notably, FAA manifests in both the light and dark cycles and possesses unique timing properties, including specific limits of entrainment and independence from known core clock components [9–11]. FAA persists even in constant darkness and in animals lacking a functional suprachiasmatic nucleus (SCN), suggesting the presence of a separate and potentially decentralized FEO network [12–15]. Unlike the relatively rigid hierarchy of light-based entrainment centered in the SCN, the system governing food anticipation appears to be highly distributed across multiple brain regions [12–14], potentially allowing for significant modulation by external factors such as metabolic state and social environment.

Historically, the vast majority of FAA studies have relied on individually housed animals, primarily due to the technical challenges of distinguishing individual behavior within a group [8, 16]. Traditional monitoring methods, such as running wheels, infrared beam breaks, or standard video tracking, lack the resolution to assign specific activity to individual animals in a shared environment [8]. While these methods have been essential in characterizing FAA, they necessarily obscure the effects of social context, which is an increasingly recognized modulator of circadian and homeostatic processes [17]. Social housing influences a wide array of physiological functions, including stress responses, metabolic profiles, and sleep architecture [18, 19], yet its influence on food-entrained rhythms remains virtually unstudied. This represents a critical gap in the field, as mice are inherently social animals, and single housing may impose an artificial state that limits ecological validity. Furthermore, while the inclusion of both sexes is becoming standard practice, the specific interaction between sex and social housing in food entrainment remains uncharacterized. We have previously demonstrated that sex and sex hormonal status significantly influence the magnitude of FAA [20, 21]. These findings suggest that males and females may employ different behavioral and physiological strategies when anticipating food, making it essential to evaluate how social context modulates these sex-specific responses.

Housing density is a primary determinant of mouse physiology, influencing metabolic rate and energy balance [17, 22]. Notably, male C57BL/6J mice exhibit increased resistance to obesity when group-housed compared to those housed individually, a phenomenon documented under both standard and high-fat diet conditions. This reduced susceptibility to weight gain and adiposity is often attributed to social thermoregulation and attenuated stress [17, 22]. Alongside these metabolic shifts, temperature rhythms serve as a sensitive marker of circadian entrainment to feeding. Mice develop robust pre-meal increases in core body temperature in response to timed feeding, a physiological response that can equal or exceed the strength of locomotor FAA [23]. Previous studies using intraperitoneal temperature monitoring have shown that these thermal profiles can distinguish between the effects of time-restricted and CR feeding [24]. Together, these findings suggest that housing conditions, sex, and internal physiological markers must be integrated to fully understand FAA under ecologically relevant conditions.

To address these gaps, we investigated FAA in male and female C57BL/6J mice, the most widely utilized inbred mouse strain in circadian research. We employed miniaturized, subcutaneously implanted wireless telemetric devices, “nanotags” [25, 26], to monitor subcutaneous body temperature and locomotor activity in both singly and group-housed animals subjected to a time-restricted feeding schedule. Crucially, this methodology enabled continuous, individual-level tracking of behavior and physiology within a socially enriched environment. Our goal was to determine whether social housing alters the timing, magnitude, or variability of FAA, and whether these effects differ by sex. To our knowledge, this is the first formal investigation of the behavioral and thermal hallmarks of FAA in socially housed mice.

## Results

### Evaluating the Effects of Cage Size on Food-Entrained Activity and Temperature Rhythms in Single Housed Male Mice

To minimize aggression among recently tagged males, animals were housed in larger enclosures (874 cm^2^; 46 × 19 × 15 cm) compared to standard small cages (468 cm^2^; 26 x 18 x 12.5 cm). Because the impact of cage volume on FAA and associated temperature rhythms is not well established, we directly compared timed restricted feeding (ZT 7-9) in male C57BL/6J mice housed in small and large cages. Food intake was comparable between housing conditions throughout the experimental period (main effect of cage size: two-way ANOVA F_1_,_18_ = 0.86, p = 0.3668; **Fig. 1A**). While body weight loss was mostly similar between groups, we observed that single housed mice had lower weight on day 14 (p < 0.0001) but not on other days (p > 0.05, **Fig. 1B**).

**Figure 1.**
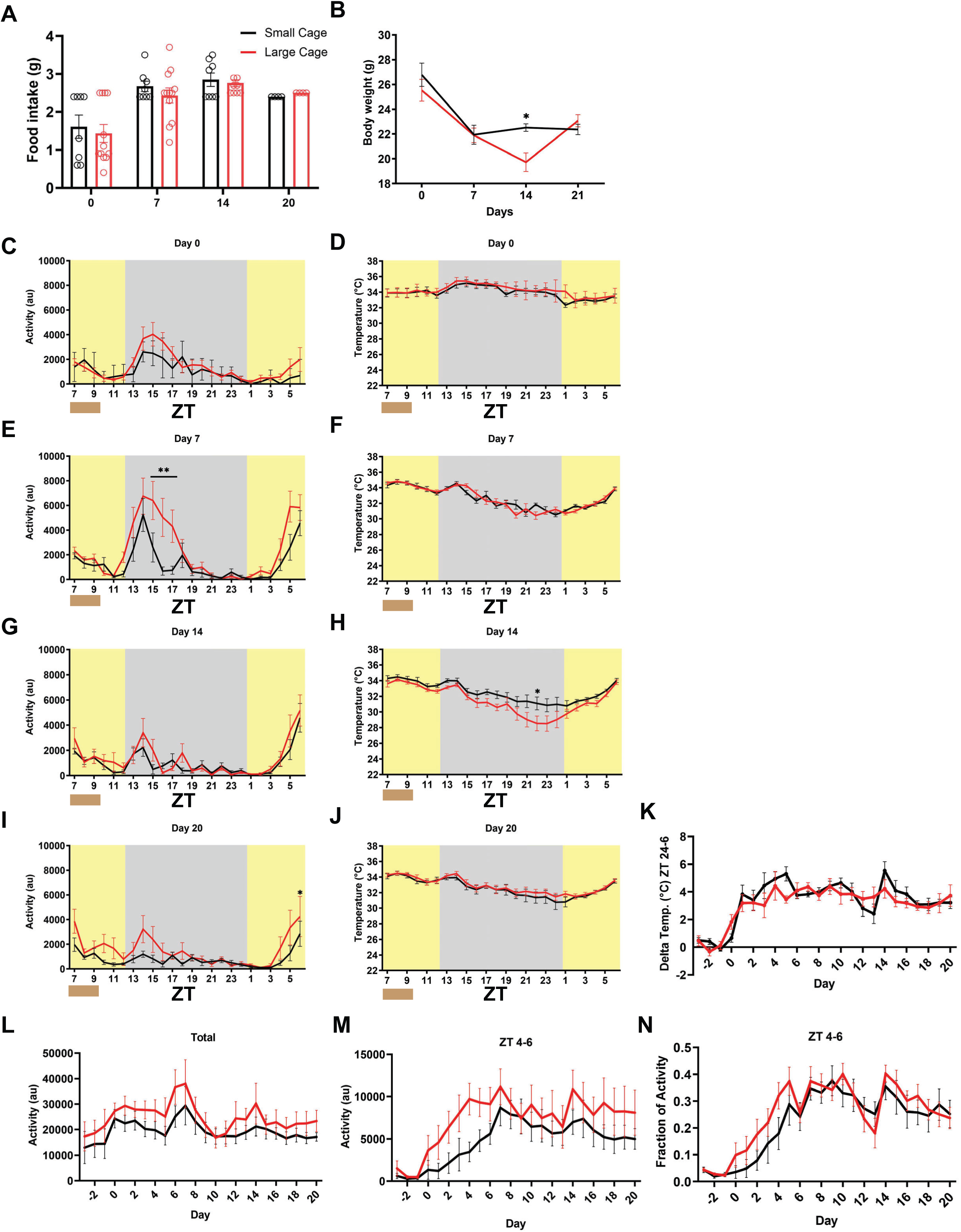
Cage size does not alter food anticipatory activity or temperature rhythms in male mice. Food intake, body weight, locomotor activity, and body temperature were compared between male C57BL/6J mice housed in small (n=8) or large (n=12) cages during restricted feeding. **(A)** Food intake across the study. **(B)** Body weight across the experiment. **(C–J)** Representative 24-h activity and temperature profiles for days 0 **(C-D)**, 7 **(E-F)**, 14 **(G-H)**, and 20 **(I-J)** of restricted feeding. Yellow shading indicates light phase; gray shading indicates dark phase; a brown line (ZT 7-9) indicates the time of food availability. **(K)** Change in body temperature (Δ temperature; ZT24–6) across days. **(L)** Total daily locomotor activity across the experiment. **(M)** Total daily locomotor pre-meal activity (ZT4–6) throughout the study. **(N)** Normalized fractional total daily activity during the pre-meal window (ZT4–6). Data are shown as mean ± SEM. Asterisk indicates a significant difference between housing conditions (*p < 0.05).. Statistical comparisons were performed using two-way ANOVA with Šídák’s multiple comparisons test where appropriate.

To assess how activity levels and subcutaneous temperature levels were affected by scheduled feeding, we monitored these variables continuously using implanted nanotags (**Fig. 1C–N**). On day 0, prior to the development of FAA, activity and temperature profiles were similar between groups (p = 0.286 for activity and p = 0.7223 for temperature; **Fig. 1C–D**). On day 7, activity showed a significant interaction of time x housing condition (F_23,414_ = 2.11, p = 0.0022), where large-caged mice exhibited a transient increase in activity during the early dark phase (ZT15–17; p < 0.05; **Fig. 1E**). However, this effect did not extend to body temperature (p = 0.218; **Fig. 1F**), and by days 14 and 20, both activity and temperature rhythms were largely identical between housing conditions at any Zeitgeber time (Šídák-adjusted post hoc tests, p > 0.39; **Fig. 1G–J**).

To quantify the development of FAA, we analyzed locomotor metrics across the experiment. Total daily activity varied significantly across days (F_23,357_ = 5.72, p < 0.0001) but was not influenced by cage size (p = 0.4966; **Fig. 1L**). As expected, pre-meal activity (ZT4–ZT6) increased as the mice entrained to the feeding schedule (F_3,216_ = 4.84, p = 0.0040), yet no significant differences emerged between small and large cages in either total pre-meal counts (p = 0.5147; **Fig. 1M**) or the fraction of daily activity occurring during the anticipatory window (p = 0.8523; **Fig. 1N**).

Daily changes in body temperature from ZT24 (the typical nadir) to ZT6 (when feeding occurs), were expressed as delta (Δ) temperature, across the course of the experiment (**Fig. 1K**). Although fluctuations in temperature change were observed across days in both housing conditions, these changes occurred in parallel, indicating that cage size did not significantly influence daily temperature responses. A mixed-effects model (repeated measures) revealed a significant main effect of time on temperature change (F_4,67_ = 12.71, p < 0.0001), indicating that there was a change in temperature change over the duration of time-restricted feeding. In contrast, there was no significant main effect of housing condition (F_1,18_ = 0.01786, p = 0.8952), nor was there a significant interaction of time and housing (F_4,67_ = 1.185, P = 0.3260), suggesting that days were similar between small-caged and large-caged mice. Post hoc Šídák’s multiple comparisons tests were used to compare temperature change between housing conditions on each day of the experiment. No individual day exhibited a statistically significant difference between groups following correction for multiple comparisons (all adjusted p ≥ 0.86). Together, these results demonstrate that the increased cage volume required for group housing does not substantially alter feeding behavior, metabolic demand, or the circadian expression of FAA and its associated thermal rhythms

### Comparison of food intake, body weight, activity, and temperature in single and group housed male C57BL/6J mice

We next examined the influence of social context on the development and expression of FAA by comparing single- and group-housed (n=4 per cage) male mice. Food intake did not differ significantly between housing conditions (main effect of housing: F_1,18_ = 1.25, p = 0.2783; **Fig. 2A**), suggesting that differences in caloric consumption do not drive the observed phenotypes. In contrast, body weight diverged significantly over time (housing x time interaction: F_3,54_ = 11.23, p < 0.0001), with group-housed mice weighing more than single-housed controls at Day 7 (p = 0.0248), Day 14 (p = 0.0011), and Day 21 (p = 0.0015; **Fig. 2B**).

**Figure 2.**
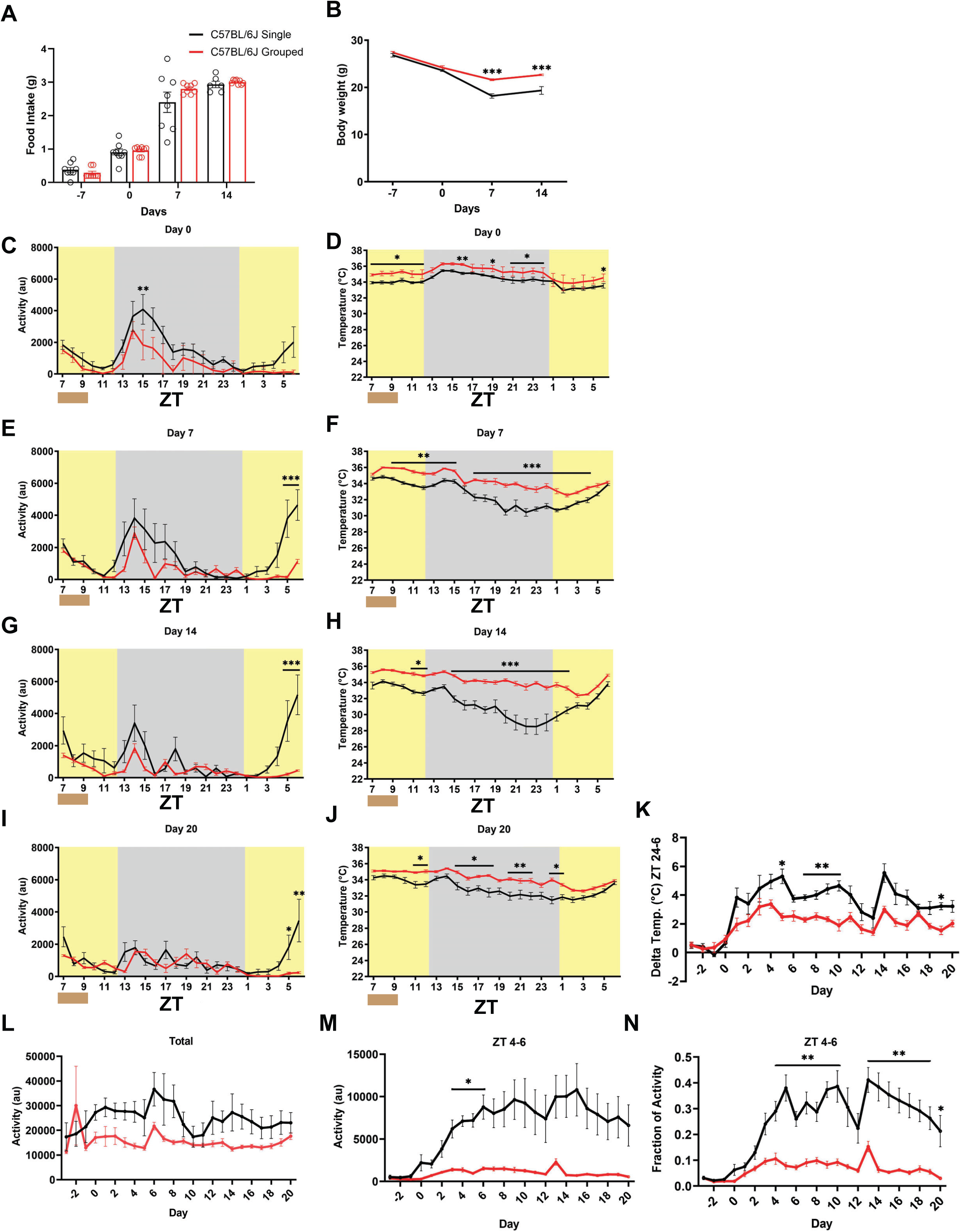
Social housing suppresses food anticipatory activity and temperature rhythms in male mice. Food intake, body weight, locomotor activity, and body temperature were compared between single (n=12) and group-housed (n=8) male C57BL/6J mice during restricted feeding. **(A)** Food intake across the experiment. **(B)** Body weight across the study. **(C–J)** Representative 24-h activity and temperature profiles on days 0 **(C-D)**, 7 **(E-F)**, 14 **(G-H)**, and 20 **(I-J)**. Yellow shading indicates the light phase; gray shading indicates the dark phase; a brown line (ZT 7-9) indicates the time of food availability. **(K)** Change in body temperature (Δ temperature; ZT24–6) across the study. **(L)** Total daily locomotor activity across the experiment. **(M)** Total daily locomotor pre-meal activity (ZT4–6) throughout the study. **(N)** Normalized fractional total daily activity during the pre-meal window (ZT4–6). Data are shown as mean ± SEM. Statistical comparisons were performed using a two-way ANOVA with Šídák’s multiple comparisons test. Asterisks indicate significant differences between housing conditions (*p < 0.05, **p < 0.01, ***p < 0.001).

Daily activity and body temperature rhythms were assessed across the course of restricted feeding (**Fig. 2C–N**). Prior to the start of the scheduled feeding (Day 0), activity levels were strongly nocturnal and temperature profiles were consistently lower in single housed mice but also showed a clear elevation at night (F_23,414_ = 1.100 and p = 0.341 and F_23,345_ = 0.8955 and p = 0.6051 respectively for activity and temperature; **Fig. 2C–D**). As restricted feeding progressed, single-housed mice developed robust FAA, whereas group-housed mice displayed a marked suppression of anticipatory behavior (Day 7 interaction: F_23,414_ = 2.334, p < 0.0005; **Fig. 2E, G, I**). This behavioral suppression was accompanied by a modest attenuation in preprandial temperature rises in the group-housed cohort (Day 7 interaction: F_23,414_ = 2.436, p = 0.0003; **Fig. 2F, H, J**).

Notably, individual thermal traces revealed a significant difference in thermoregulatory stability (**Supp. Fig. 1**). While 8 out of 8 group-housed male mice maintained stable temperatures, 6 out of 12 of the single-housed males exhibited acute hypothermic bouts (subcutaneous temperature < 28°C) during the mid-light phase (p = 0.0419, Fisher’s exact test; **Supp. Fig. 1**).

To quantify these adaptations, we analyzed summary metrics across the duration of the experiment. Total daily locomotor activity was equivalent between groups (F_1,18_ = 1.63, p = 0.2181; **Fig. 2L**), indicating that the suppression of FAA was not a result of generalized lethargy. However, both pre-meal activity (ZT4–6; F_1,18_ = 13.91, p = 0.0015; **Fig. 2M**) and the fraction of daily activity occurring during the anticipatory window (F_1,18_ = 11.66, p = 0.0031; **Fig. 2N**) were significantly reduced in group-housed mice.

Finally, the daily delta temperature (ZT 24 to ZT 6 change) was slightly lower in the social housing condition (F_1,18_ = 25.67, p < 0.0001; **Fig. 2N**). For the daily ZT24–ZT6 temperature change in male mice, a mixed-effects model revealed a significant effect of time (F_4,79_ = 14.98, p < 0.0001), and a main effect of housing condition (F_1,18_ = 25.67, p < 0.0001) and no effect of the time × housing interaction (F_4,79_) = 2.10, p = 0.0773; Figure 2K). Single-housed males showed a numerically greater mean temperature change than group-housed males (3.394°C vs. 1.974°C; mean difference = 1.419°C; 95% CI = 0.8307 to 2.008), indicating a significant overall difference between conditions. Consistent with this, post hoc Šídák’s multiple-comparisons tests identified significant differences between single- and group-housed males on certain days (days 5, 7-10, and 19; adjusted p ≤ 0.0276), while no differences were detected on other days. Together, these findings demonstrate that social housing robustly suppresses the magnitude of FAA and its associated thermal rhythms without altering overall daily activity or food intake during restriction.

### Comparison of food intake, body weight, activity, and temperature in single and group housed female C57BL/6J mice

We next examined the impact of social housing on female C57BL/6J mice. Food intake was largely similar between groups, though single-housed females exhibited a transient increase in consumption at Day 7 (housing x time interaction: F_3,24_ = 15.66, p < 0.0001; **Fig. 3A**). Post hoc analysis confirmed that single-housed mice consumed significantly more food on Day 7 (p < 0.0001) but were otherwise indistinguishable from group-housed mice on Days 14 and 21 (p > 0.9999). Unlike males, female body weight did not differ significantly by housing condition throughout the experiment (main effect of housing: p = 0.1573; **Fig. 3B**).

**Figure 3.**
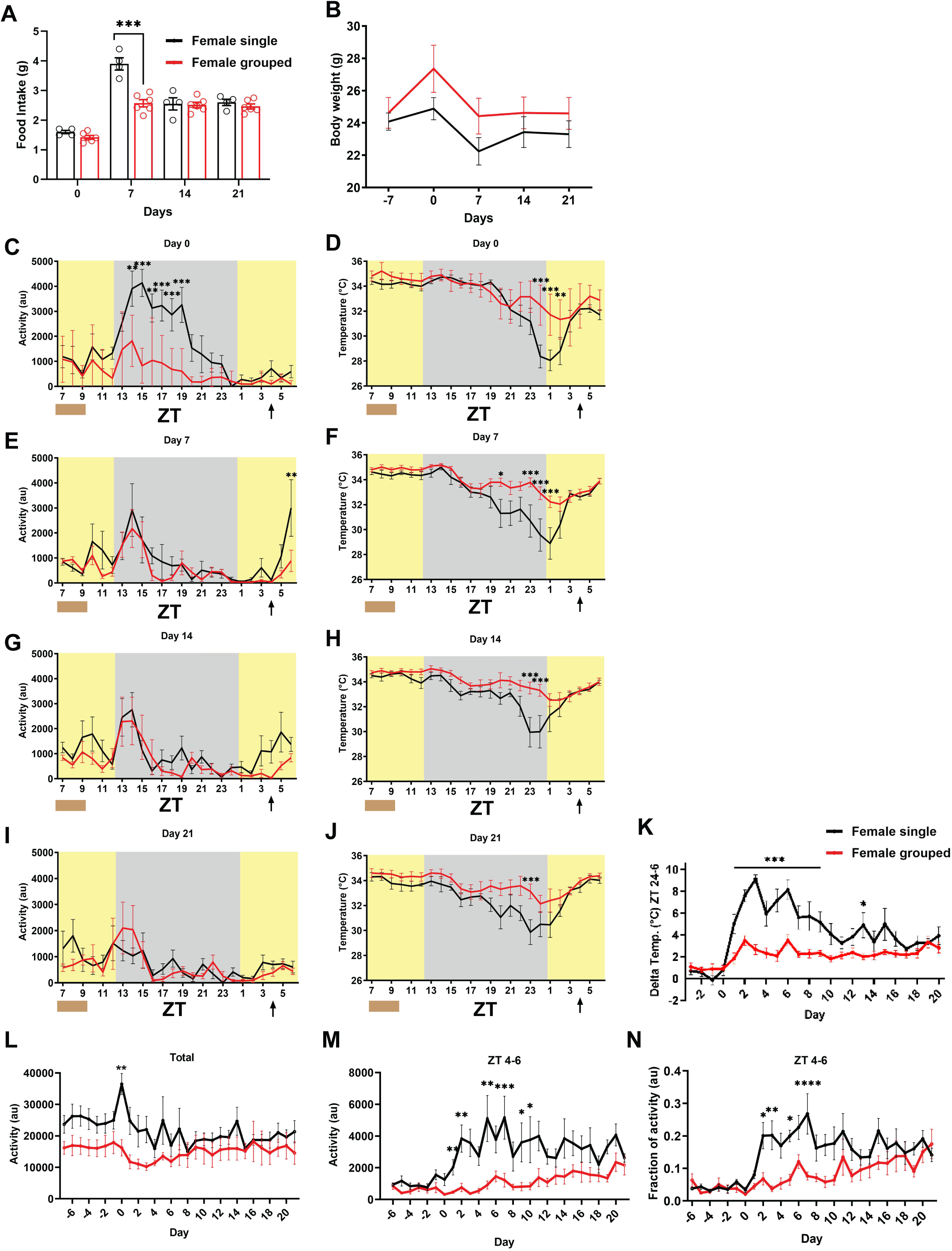
Social housing has a transient and attenuated effect on food anticipatory activity in female mice. Food intake, body weight, locomotor activity, and body temperature were compared between single (n=7) and group-housed (n=8) female C57BL/6J mice during restricted feeding. **(A)** Food intake across the experiment. **(B)** Body weight across the study. **(C–J)** Representative 24-h activity and temperature profiles on days 0 (C-D), 7 (E-F), 14 (G-H), and 21 (I-J). Yellow shading indicates the light phase; gray shading indicates the dark phase; a brown line (ZT 7-9) indicates the time of food availability. **(K)** Change in body temperature (Δ temperature; ZT24–6) across days. **(L)** Total daily locomotor activity across the experiment. **(M)** Total daily locomotor pre-meal activity (ZT4–6) throughout the study. **(N)** Normalized fractional total daily activity during the pre-meal window (ZT4–6). ( Data are shown as mean ± SEM. Statistical comparisons were performed using two-way ANOVA with Šídák’s multiple comparisons test. Asterisks indicate significant differences between housing conditions.

Daily activity and body temperature rhythms were monitored continuously and plotted for Days 0, 7, 14, and 21 (**Fig. 3C–N**). On Day 0, single-housed females exhibited higher nocturnal activity amplitude compared to group-housed mice (p < 0.002; **Fig. 3C**). As restricted feeding progressed, single-housed females developed robust FAA by Day 7, whereas group-housed females showed a significantly weaker and delayed anticipatory response (**Fig. 3E**). However, these differences diminished over time; by Day 14, FAA magnitude began to converge (**Fig. 3G**), and by Day 21, activity and temperature profiles were largely similar between housing conditions (**Fig. 3I–J**). To quantify these transient effects, we analyzed summary metrics across the experiment. Total daily locomotor activity was significantly higher in single-housed females only on Day 0 (p = 0.0012) and was equivalent between groups thereafter (**Fig. 3L**). However, **pre-meal activity** (ZT4–6) was significantly greater in single-housed females during the initial week of restricted feeding (Days 2, 3, 5, and 7; p < 0.05), with no significant differences observed in the later stages of the study (**Fig. 3M**). We also calculated the fraction of daily activity occurring during the pre-meal window to account for individual differences in baseline arousal. This fractional FAA was significantly higher in single-housed females on Days 1, 3, 5, and 7 (p < 0.05), before converging with the group-housed cohort for the remainder of the experiment (**Fig. 3N**).

The daily ZT24–ZT6 temperature change was analyzed in female mice with a two-way repeated-measures ANOVA (**Fig. 3K**). This test revealed significant effects of day (F_23,299_ = 14.57, p < 0.0001), housing condition (F_1,13_ = 20.96, p = 0.0005), and a significant day × housing interaction (F_23,299_ = 6.092, p < 0.0001). Group-housed females exhibited a lower mean temperature change than single-housed females (2.174°C vs. 4.270°C; mean difference = −2.096°C; 95% CI = −3.085 to −1.107), a noticeable effect of the housing condition. Post hoc Šídák’s multiple-comparisons tests showed that single-housed females had significantly higher temperature changes than group-housed females on multiple consecutive days, specifically from day 1 through day 9 (adjusted p values ranging from 0.0090 to <0.0001), as well as on day 13 (p = 0.0330). No differences were detected at baseline or on later time points (all adjusted p ≥ 0.1061). Together, these results indicate that social housing exerts a more modest and transient influence on FAA in female mice compared to males, primarily affecting the early adaptation to restricted feeding.

## Discussion

Using implanted sensors to obtain individual-level activity and body temperature measurements in group-housed animals, we show that social housing robustly attenuates FAA and blunts preprandial increases in body temperature. This suppression was observed in both sexes, with the strongest effect in male mice. These findings identify social context as a powerful and underappreciated modulator of food-entrained circadian behavior.

FAA has traditionally been viewed as a robust circadian phenomenon driven by a FEO operating independently of the light-entrained suprachiasmatic nucleus [5, 6, 9]. However, nearly all prior studies examining FAA have been conducted in singly housed animals implicitly assuming that FAA magnitude and expression are invariant across social environments [8]. Our results challenge this assumption. Group housing substantially reduced anticipatory locomotor activity without abolishing it, indicating that while food-based entrainment persists, its behavioral expression is strongly shaped by group housing.

The interpretation of these findings is informed by prior work demonstrating that locomotor and thermal components of FAA can be differentially regulated. For example, preprandial increases in body temperature can be observed under conditions where locomotor anticipatory activity is reduced or absent, and conversely, behavioral anticipation can occur with attenuated temperature rhythms [23, 27, 28]. These observations indicate that FAA is not an output of a single oscillator, but instead reflects the coordinated activity of multiple food-entrainable oscillators that regulate distinct physiological and behavioral processes. In this context, the parallel suppression of both activity and temperature observed under social housing suggests a coordinated shift in underlying physiological state rather than a simple attenuation of motor output.

An insight from this study is that the effects of social housing on anticipatory behavior are paralleled by changes in body temperature rhythms. Single-housed mice exhibited robust preprandial increases in temperature, whereas group-housed mice showed markedly attenuated temperature responses. Because body temperature is tightly coupled with metabolic rate and energy expenditure [29–31], these findings suggest that FAA is not solely a behavioral readout of circadian timing but also reflects underlying physiological state. Consistent with this, preprandial increases in body temperature are a well-established feature of FAA [6, 8, 9]. One plausible mechanism linking social housing to these effects is altered thermoregulatory demand. Standard laboratory temperatures (∼20–24 °C) fall below the murine thermoneutral zone (∼29–32 °C), placing singly housed mice under chronic cold stress [31, 32]. Under these conditions, animals must increase metabolic heat production to maintain core temperature [33]. In contrast, group-housed mice benefit from social thermoregulation, including huddling, which reduces heat loss and lowers energetic demand [34, 35]. Although differences in cage air temperature are modest, the effective thermal environment experienced by the animal differs substantially, as reflected in changes in core body temperature and heat conservation [36].

In this context, preprandial increases in activity and temperature may represent coordinated components of an anticipatory metabolic state that mobilizes energy in advance of food availability. When energetic demand is high, as in singly housed mice, this anticipatory response is pronounced. When energetic demand is reduced, as in group-housed mice, both activity and temperature responses are attenuated. The close alignment between locomotor and temperature changes observed here supports the idea that FAA reflects an integrated physiological state rather than a purely circadian motor output. This interpretation is further supported by prior work demonstrating that increasing ambient temperature toward thermoneutral conditions similarly reduces FAA. In singly housed mice, elevating housing temperature to ∼30 °C attenuates anticipatory locomotor activity [27], indicating that reduced thermoregulatory demand alone is sufficient to dampen FAA. Together with the present findings, these results support a model in which both environmental temperature and social context likely converge on a common energetic mechanism regulating the magnitude of anticipatory behavior. This energetic interpretation is also consistent with the observation that group-housed males maintained greater body weight than singly housed males despite similar reported food intake. One possibility is that social housing reduced thermogenic costs through improved heat conservation, allowing more ingested energy to be retained for growth or body mass maintenance rather than expended on cold-induced heat production. Consistent with this interpretation, hypothermic bouts were observed in approximately half of singly housed mice but were not detected in group-housed mice, suggesting that social housing may improve thermoregulatory stability under standard vivarium conditions. Notably, singly housed mice that experienced hypothermic bouts often showed altered activity patterns relative to mice that maintained internal temperatures above 28 °C. However, the effects of hypothermia on FAA were not directly tested here and remain outside the scope of the present study. Therefore, while these observations support a link between housing condition, thermal stability, and anticipatory behavior, future studies will be required to determine whether hypothermia contributes causally to the regulation or expression of FAA.

Differences between males and females were also apparent in the magnitude and time course of FAA suppression. Male C57BL/6J mice exhibited a more pronounced and sustained attenuation under group housing, whereas females showed a more modest and transient reduction during the early phase of restricted feeding. Importantly, we did not directly compare sexes in this study, and these differences should be interpreted cautiously. Female mice typically exhibit lower baseline levels of FAA than males [20, 21], which may limit the dynamic range available for social modulation. Consistent with this, the effects of housing in females were detectable but less robust. These observations suggest that sex may influence the sensitivity of FAA to environmental context, although further work will be required to directly test this possibility.

Several limitations should be considered. While our study focused on locomotor activity and body temperature, additional measures of metabolic rate, hormonal signaling, and neural activity within food-entrainable circuits will be required to define underlying mechanisms. In addition, group housing and cage density may have complex effects on stress levels and social hierarchies which could further influence FAA and warrant systemic investigation. All experiments were conducted under standard vivarium temperatures (∼22 °C), consistent with most studies; however, thermoneutral housing substantially alters activity, feeding behavior, and social thermoregulation. Thus, these findings should be interpreted within the context of typical laboratory conditions, where social and thermal factors jointly shape behavior.

Taken together, these findings show that social housing dampens FAA and associated temperature rhythms in mice, revealing this behavior as a socially and energetically modulated output rather than a fixed circadian program. These findings highlight the importance of considering social context when interpreting food-entrained circadian rhythms.

## Methods

### Ethics statement

The experiments described herein were approved by the California State Polytechnic, Pomona Institutional Animal Care and Use Committee (IACUC) under protocol 23.009.

### Mouse strains and husbandry

C57BL/6J mice (Jackson Laboratory, stock #000664) were used for all experiments. Male and female mice were 11–14 weeks of age at the start of restricted feeding. Animals were housed in a temperature-controlled vivarium (∼22 °C) under a 12:12 h light-dark cycle (lights on at zeitgeber time [ZT] 0, lights off at ZT12) with ad libitum access to water.

Following recovery from surgery (see below), mice were assigned to either single housing or group housing conditions. Each group-housed cage contained four mice (two implanted and two non-implanted controls). Group-housed mice were maintained in larger cages (46 × 19 × 15 cm) with four mice per cage as were singly housed mice described in Figures 2 and 3. For the mice described in Figure 1, we compared single housed mice in standard cages (26 x 18 x 12.5 cm) versus large cages. All mice were acclimated to their housing conditions for approximately two weeks prior to the start of restricted feeding.

Food was provided daily for a 3-hour window from ZT6 to ZT9 and was otherwise unavailable. Day 0 corresponds to the first day of food restriction. Mice were monitored throughout the restricted feeding period, and food intake was measured weekly. For group-housed cages, intake was measured at the cage level and normalized to the number of animals in the cage to provide per-animal estimates. Body weight was recorded weekly across the experiment. For group-housed conditions, four mice were housed per cage. Nanotag recordings were used to monitor activity and body temperature continuously across the experiment with a sampling rate of 5 minutes.

### Nanotag implantation surgery

Nanotags were obtained from Stoelting and decontaminated with ethylene oxide gas. Mice were anesthetized using isoflurane anesthesia and then a nanotag was implanted subcutaneously through a 2-cm incision on the dorsal midline. The incision was stapled using 9mm EZ Clip Wound Closure System (Braintree Scientific). We applied 0.1 mL topical anesthetic (Hospira, Marcaine 0.5% 5 mg/mL, PAA113011) to the incision site. Animals were monitored for a minimum of three days following surgery and were given 0.1mL 0.5mg/mL ketoprofen during this period. Once implanted, incisions were stapled, and the mice were returned to individual housing for recovery, receiving perioperative care in accordance with institutional guidelines. Once healed, the mice were assigned to either group housing or single housing (control).

At least two weeks post-surgery, mice were assigned to group of single conditions. For group-housed experiments, each cage contained four age-matched mice: two implanted mice used for nanotag-based behavioral and temperature monitoring and two non-implanted cage mates. This cage composition was used to maintain a standard group-housing environment while allowing individual activity and temperature measurements from identified implanted animals. No data were collected regarding the non-implanted mice.

Temperature data were acquired in 5-minute bins and aggregated for analysis across hours and days. For hourly temperature profiles, values were averaged within each hour and plotted across a 24-hour cycle spanning ZT7 to ZT6 of the following day. For daily delta temperature calculations, the maximum temperature value within the ZT6 hour (derived from 5-minute bins) and the minimum temperature value within the ZT24 hour were identified for each animal on each day. The difference between these values (maximum at ZT6 minus minimum at ZT24) was computed to yield a single delta temperature value per day, which was plotted across the included days of the experiment. In this manner, we captured the nadir and peak of temperature for each animal. At the end of the experiment, mice were euthanized by CO_2_ narcosis.

### Statistical testing

All statistical analyses were performed using GraphPad Prism (version 10.0; GraphPad Software, RRID: SCR_002798). Data are presented as mean ± SEM. Statistical significance was defined a priori as p < 0.05. For comparisons involving repeated measurements across time (e.g., body weight, food intake, locomotor activity, and temperature across days or Zeitgeber time), two-way ANOVA or mixed-effects models (restricted maximum likelihood) were used, as appropriate. Mixed-effects models were applied in cases of missing values or unequal sample sizes. Significant main effects or interactions were followed by Šídák’s multiple comparisons tests. For analyses of circadian profiles (e.g., 24-hour activity and temperature traces), two-way ANOVA with factors of time and housing condition (or cage size) was used. For single time point comparisons between two groups, unpaired two-tailed student’s t-tests were used. Welch’s correction was applied when variances were unequal. For categorical data (e.g., incidence of hypothermic events), Fisher’s exact test was used. Repeated-measures two-way ANOVA was used for longitudinal datasets. For group-housed conditions, individual animals were treated as the unit of analysis, as nanotag recordings enabled continuous, individual-level measurements independent of cage-level aggregation.

---

**Figure S1. Individual variability in temperature and activity in single- and group-housed male mice.** Individual-level activity and body temperature traces from representative mice during restricted feeding. **(A–C)** Mean temperature and activity profiles for single- and group-housed mice. **(D–N)** Representative individual traces showing activity (solid line) and body temperature (dashed line) across the light-dark cycle. Single-housed mice exhibited substantial heterogeneity in temperature regulation, with a subset displaying pronounced reductions in body temperature during the mid-light phase across restricted feeding days. These hypothermic episodes were not observed in group-housed mice, which exhibited more stable temperature profiles. Yellow shading indicates the light phase; gray shading indicates the dark phase; a brown line (ZT 7-9) indicates the time of food availability.

Supporting Information: Minimal data set. Excel file containing data used to prepare the Figures.

