## Supplementary figures and images for "Social Context Suppresses Food Anticipatory Activity and Associated Thermoregulation in Mice"

### Supplemental Figure 1

#### Supplemental Figure 1

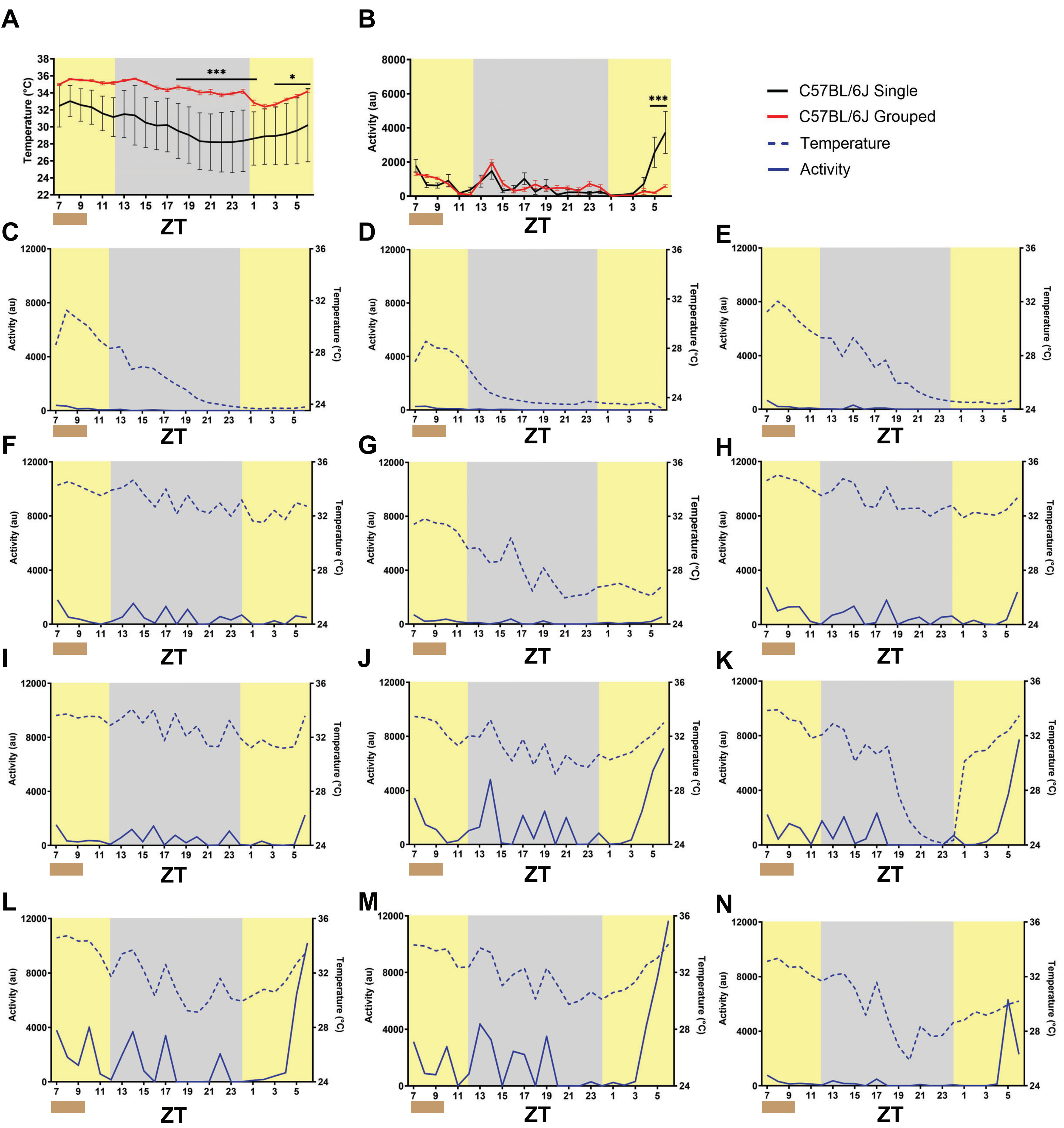
